# Estimating butterfly population trends from sparse monitoring data using Generalized Additive Models

**DOI:** 10.1101/2023.12.07.570644

**Authors:** Collin Edwards, Cheryl Schultz, David Sinclair, Daniel Marschalek, Elizabeth Crone

## Abstract

Concerns of declines in insects and population level responses to climate change have highlighted the importance of estimating trends in abundance and phenology from existing monitoring data. As the taxa with the most systematic monitoring data, butterflies are a frequent focus for understanding trends in insects. Even so, ecologists often have only sparse monitoring data for at-risk butterfly populations. As existing statistical techniques are typically poorly suited to such data, these at-risk populations are frequently excluded from analyses of butterfly trends. Here we present guidelines for estimating population trends from sparse butterfly monitoring data using generalized additive models (GAMs), based on extensive simulations and our experiences fitting hundreds of butterfly species. These recommendations include pre-processing steps, model structure choices, and post-hoc analysis decisions that reduce bias and prevent or mitigate biologically implausible model fits. We also present the ButterflyGamSim package for the programming language R, available at https://github.com/cbedwards/butterflyGamSims. This open source software provides tools for ecologists and applied statisticians to simulate realistic butterfly monitoring data and test the efficacy of different GAM model choices or monitoring schemes.

---

A key task in both population ecology and conservation biology is to infer population trends – both growth rates and, increasingly, trends in phenology – from monitoring data. These trends are the foundations for studies in basic ecology, study of species of conservation concerns (e.g., Bonoan et al. 2021), and comparative studies identifying overarching patterns in abundance and phenology across species (Diamond et al. 2011, Forister et al. 2021). Many species, such as butterflies, have short activity periods during which they are conspicuous, with the timing of peak activity varying from year to year. Monitoring programs for such species are often structured around repeated surveys across a year, in turn repeated across years and sometimes across sites (e.g. Pollard Walk design) (Pollard 1994, Shapiro 2020, PollardBase). Ecologists thus need practical tools for estimating yearly quantities of abundance and phenology, which can then be scaled up to identify variation and trends across years.

Many tools have been developed to translate repeated yearly surveys into yearly estimates of population characteristics (Edwards and Crone 2021), particularly for butterflies. Fundamental to most of these tools are underlying assumptions about the shape of activity: some methods assume survey counts can be approximated with a unimodal, gaussian shape across a single year (Lindén and Mäntyniemi 2011, Dennis et al. 2015, Stewart et al. 2020, Edwards and Crone 2021); the Zonneveld model and the epsilon-skew-normal models fit unimodal shapes with varying degrees of skew and kurtosis (Zonneveld 1991, Clark and Thompson 2011); Gaussian mixture models can represent multimodal activities as might be expected for multivoltine species, presuming activity can be decomposed into Gaussian curves (Proïa et al. 2016). In contrast to most methods, general additive models (GAMs) in the form of smoothing splines make very few assumptions about the shape of activity, making them a strong choice (a) for analyses when aspects of yearly activity are not known ahead of time and must be inferred from the data (i.e., uncertain or shifting voltinism), or (b) to provide a consistent framework for comparative analysis including species with diverse patterns of activity (Rothery and Roy 2001, Hodgson et al. 2011, Stemkovski et al. 2020).

The key benefit GAMs provide is the ability to specify a predictor (e.g., day of year) as having some unknown, potentially nonlinear relationship with the response (e.g., butterflies seen). Following Pederson et al. (2019), we will refer to the terms representing these relationships as “smoothers”. Because these smoothers are flexible, GAMs are able to allow data to identify the relationship (including any nonlinearity) between a predictor day of year and butterfly count, rather than having that relationship be dictated by the modeler. However, the flexibility of smoothing splines leaves them prone to overfitting when working with sparse data, as is common for some types of population monitoring data. The simple solution of dropping years or populations with limited data will bias inferences made from estimated trends (Didham et al. 2020), particularly since populations with more limited data have been found to be disproportionately at-risk species (Forister et al. 2023). Here we offer specific approaches – pre-processing steps, model structures, and post-hoc analysis steps – that perform well for sparse monitoring data. For those looking to understand smoothing splines and how to use them, we highly recommend Noam Ross’s free interactive course “GAMs in R” (https://noamross.github.io/gams-in-r-course/).

To our knowledge, there are no consistent guidelines for using smoothing splines to estimate population trends from monitoring data, particularly for sparse monitoring data. We present guidelines and approaches we have developed in the process of fitting monitoring data for hundreds of butterfly species, informed by extensive simulation using our ‘butterflyGamSim’ R package. These guidelines are targeted for working with sparse butterfly monitoring data from temperate regions (with seasonal rather than year-round activity). However, many of our considerations are relevant for other systems (e.g. choosing appropriate model structure, post-hoc analysis for populations that appear to go extinct). We provide implementation examples specifically for the package ‘mgcv’ (Wood 2011) in the R programming language (R Development Core Team 2023), but our guidelines are relevant for other software implementations of smoothing splines.

We begin with an overview of using smoothing splines, describe our new butterflyGamSims package, and then provide our recommendations. Simulation results are summarized in the supplements; all simulation results will be made available upon publication.

## Smoothing spline method overview

Like many statistical approaches for estimating trends from monitoring data, we first fit a model to the data, then extract metrics of interest (i.e., abundance index, phenology metrics) from the fitted model (Clark and Thompson 2011, Hodgson et al. 2011, Edwards and Crone 2021) (Fig. 1A-B). For some other types of models, yearly metrics of interest can be identified analytically from parameter estimates (e.g., Gaussian, Zonneveld, epsilon-skew-normal models). Because smoothing splines are semi-parametric and their parameters are less obviously linked to metrics of interest, we obtain our yearly metrics numerically. To do so, we use the fitted model to predict activity for each year (“activity curves”) and calculate metrics from these predictions for each year (Fig. 1C). These yearly metrics act as single data points per year, which can then be analyzed for across-year trends (Fig. 1D), and uncertainty in those trends can be estimated using non-parametric bootstrapping (Efron 1992). We note that some other modeling approaches can allow the initial model to directly incorporate parameters for trends in abundance or phenology (e.g., Macphie et al. 2023), but presently there is no easy way to do so with smoothing splines while allowing the activity curve to change shape or timing between years.

**Figure 1:**
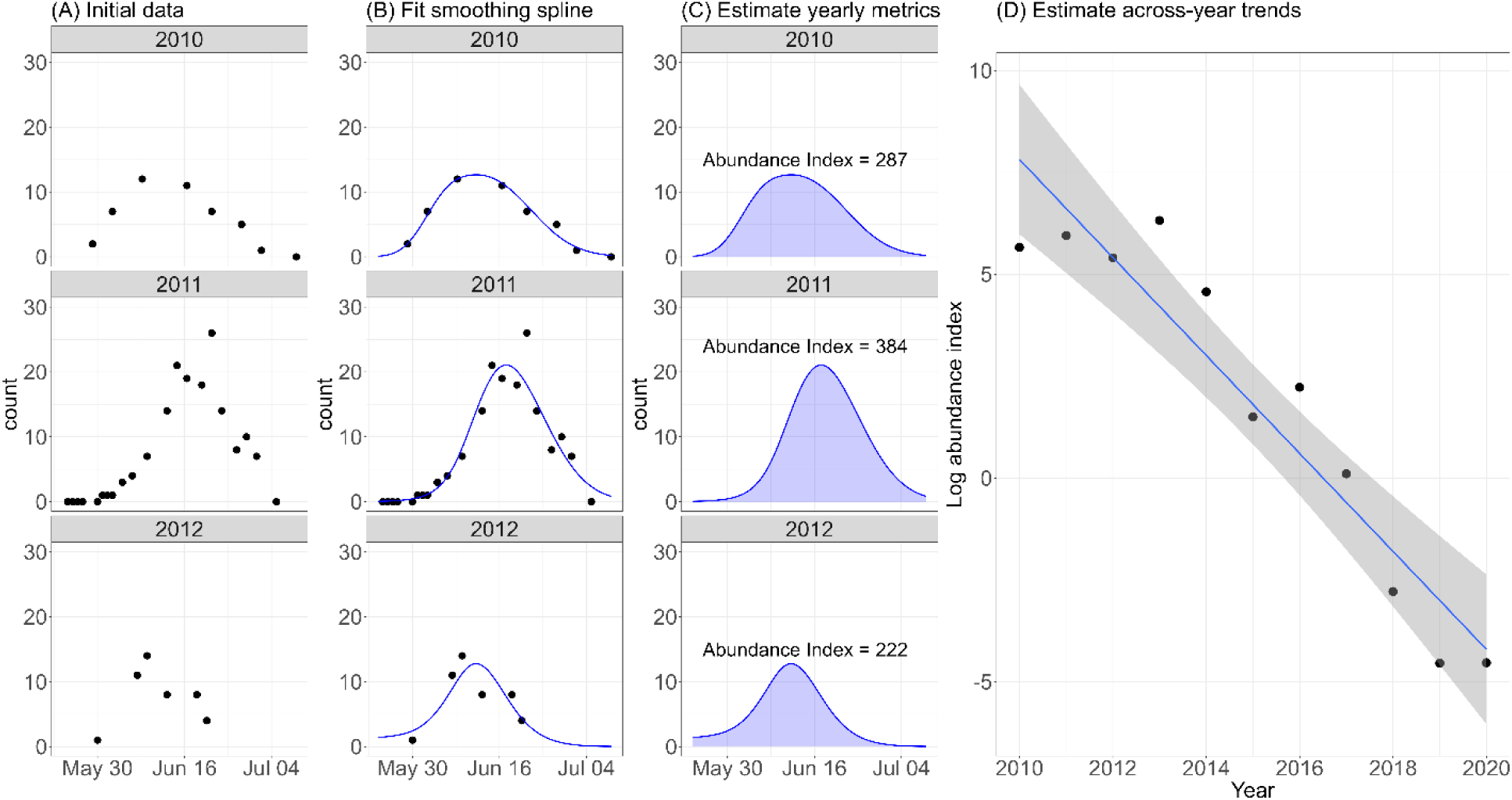
Overview of estimating population trends using smoothing splines, using data from monitoring a population of the Karner blue butterfly. Starting from monitoring data with repeated sampling within years (A), we fit smoothing spline to the data, and use the prediction of the fitted model to produce the activity curves (B). Various yearly population metrics can be estimated from the predictions, including abundance index (calculated as the area under the curve) (C). From these yearly metrics, trends across years can be estimated (D), including growth rate (trend in log abundance index across time).

We focus on the previously published yearly metrics of abundance index, median phenology, onset phenology, end of activity, and flight period (i.e., Bonoan et al. 2021 Figure 2). Abundance index for a year is the area under the activity curve, and is a measure of observable-activity-days, analogous to the simpler and coarser practice of estimating abundance as the sum of counts across all surveys in a year (e.g., Pollard 1977). Under the assumption that there is no systematic change in observability across years, growth rate can be estimated as the slope of log abundance index across years. For phenology metrics, we follow Miller Rushing et al. (2008), van Strien et al. (2008) and Inouye et al. (2019) in strongly advising the use of quantile-based metrics. Metrics based on the absolute predicted count (e.g., the day butterflies are first observed, or the first day the activity curve equals or exceeds 1) will confound phenology with abundance and create an apparent tendency for years of higher abundance to have earlier starts and later ends of season, which is usually not a desirable trait (but see Bonoan et al. 2021). This is because with more individuals there are higher chances of at least one individual having unusually early or late activity, but this does not reflect changes in the expected phenology of any given individual. Like others, we find that onset and end of activity are reasonably represented by the 0.1 and 0.9 quantile of the activity curves (Jonzen et al. 2006, Michielini et al. 2020), although other quantiles can also be used (e.g., Linden et al. 2017). Median phenology (the 0.5 quantile) serves as a reasonable measure of the central timing of activity, and is robust to multimodal distributions in ways that day of peak abundance is not (but see “cautions” and Fig. 7 for additional considerations with bivoltine species). We calculate the flight period as the number of days between onset and end, i.e., the length in days of the middle 80% of activity.

**Figure 2:**
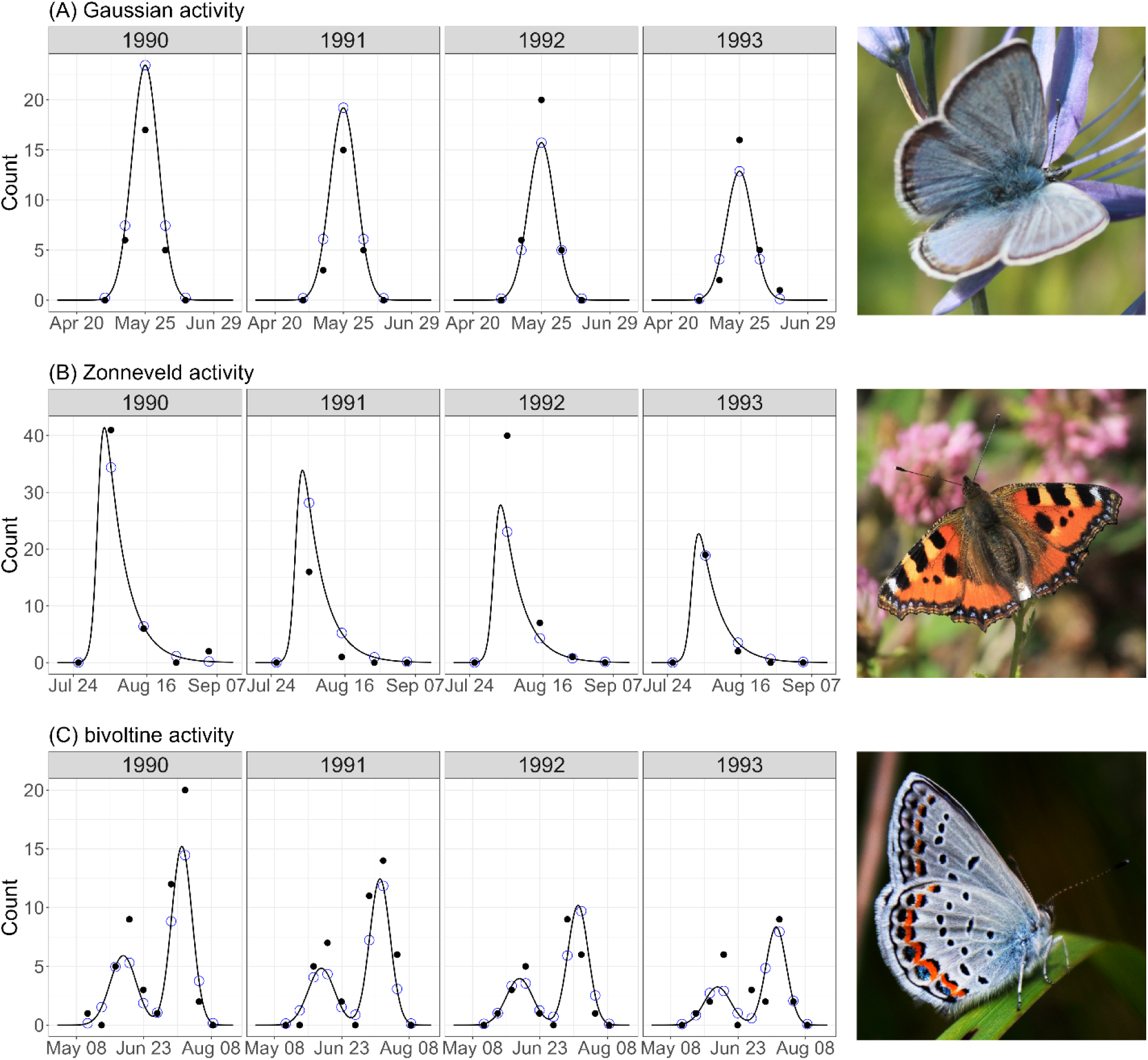
Example timeseries generated from ButterflyGamSim package for three qualitatively different activity curves: (A) Gaussian activity curves parameterized for the Fender’s blue butterfly, (B) Zonneveld activity curves parameterized for the small tortoiseshell butterfly, and (C) bivoltine activity curves parameterized for the Karner blue butterfly. Black curves show the underlying activity curve representing the “real” simulated biology; blue circles show the expected “real” activity on days of simulated sampling; black points show the counts of simulated monitoring events, based on the expected activity but incorporating sampling error. (D) Fender’s blue butterfly, (E) small tortoiseshell butterfly, (F) Karner blue butterfly. [Photo credits: (D) Jani Uusitalo, (E) Aaron Carlson, (F) C.S.]

## The butterflyGamSims package

To identify best practices for fitting sparse butterfly monitoring data, we developed a simulation framework to generate realistic monitoring data with known underlying population behavior and then estimate population trends using specified model structures and fitting approaches. In this framework, we specify the shape of the activity curve and a growth rate for the population. For each year, the simulation first determines the relative population size of that year based on the previous year and growth rate, then calculates the corresponding activity curve. Finally the simulation generates simulated monitoring counts on user-specified days, using the height of the activity curve on those days and a discrete probability distribution (e.g. Poisson, negative binomial) to generate count data that includes sampling variability. Using this framework, we simulated 1000 20- and 30-year time series of butterfly population monitoring data for each of three qualitatively distinct patterns of yearly butterfly activity: gaussian activity based on parameterization of Fender’s Blue butterflies estimated in Bonoan et al. (2021); Zonneveld-shaped activity based on parameterization of the small tortoiseshell butterfly estimated in Zonneveld (1991); and bivoltine activity based on our fitting of Karner Blue butterfly (Fig. 2).

Our focus was identifying methods that worked even on sparse data sets that are sometimes removed from broader analyses. Therefore, based on our experience with sparse monitoring data, we simulated only 5 days of monitoring per butterfly generation per year, evenly distributed to span the period of activity. Because we have observed that smoothing splines perform particularly poorly on time series when initial or final years consist mostly of zero counts, we simulated two sets of population dynamics: (a) 20-year time series of strong population declines (growth.rate = −0.12) with initial population size chosen such that simulated data frequently had mostly zero observations in the final years of the time series (init.size = 1000), and (b) 30-year time series of even more extreme declines (growth rate = −.18) with smaller initial population sizes (400) chosen such that simulated data frequently ended with several years of all zero-counts. Additional simulations using less extreme growth rates (−0.06) showed qualitative agreement with our findings, but model fits and inferences were less problematic even without our various suggested best practices. Smoothing splines and post-hoc analyses behave identically when reversing the order of years, so methods that worked well for our simulated data would work equally well for populations starting at very low abundance and increasing with growth rates of 0.12 or 0.18. For this reason we did not also simulate growing populations. To simplify interpretation of results, we simulated static phenology. Thus, when comparing across simulations, unbiased methods would have estimated growth rates centered on −0.12 or −0.18 and estimated phenology trends centered on 0. The full list of simulation parameters used can be found in Table 1.

**Table 1:**
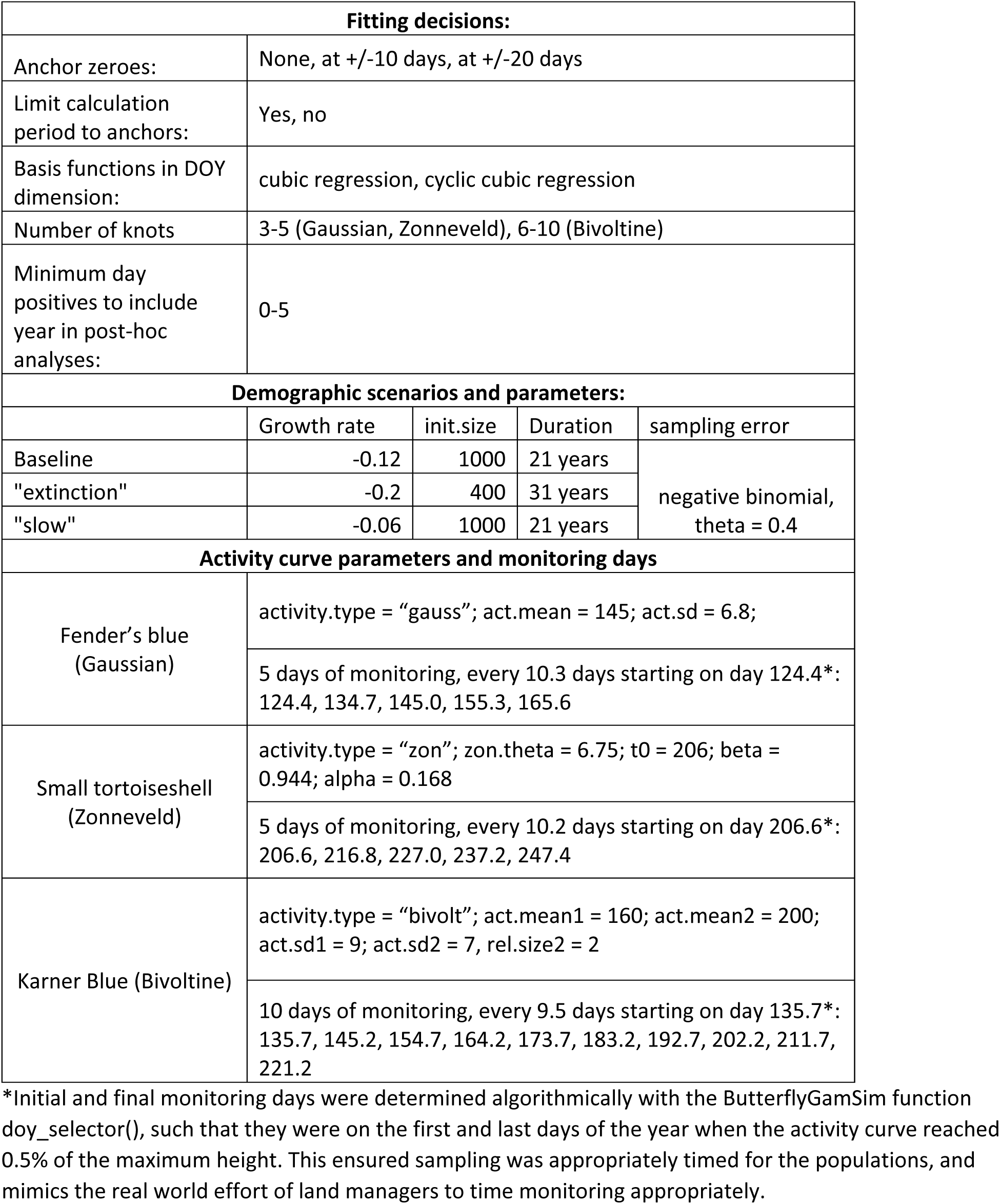
Parameter values used in simulations.

Using our simulated data, we tested the efficacy of a range of modeling choices, which are described in later sections. These included: (a) using no anchor zeroes (see “avoiding misfits”, below), anchor zeroes at 10 days, or anchor zeroes at 20 days, (b) using cubic regression or cyclic cubic regression smoothers in the day of year (DOY) dimensions, (c) limiting our estimation of yearly metrics to only the period between the data or using the entire year, (d) using 3-5 knots in the DOY dimension for univoltine simulations or 6-10 knots for the bivoltine simulations, and (e) excluding yearly estimates from post-hoc analysis if the number of day-positives (monitoring days with one or more butterfly seen) was below a threshold, with a threshold varying from 0 to 5 (Table 2). To summarize the efficacy of different fitting choices, for each combination of fitting options and for each type of trend (growth rate, phenology trends) we compared the estimates to the specified true trends. An ideal method would have little variation between simulations (low variance), and the average estimated trend across simulations would be close to the true trend (low bias) (James et al. 2021). When applying the same analysis approaches to three distinct types of simulated activity curves (Gaussian, Zonneveld, bivoltine), sometimes different fitting choices were optimal for different types of activity curve. Since in practice we are unlikely to know the exact shape of activity for our data (aside from sometimes knowing the number of generations), we focus on modeling decisions that are consistently optimal across the three types of simulated data, and explicitly state when modeling choices have less clear “best” choices.

**Table 2:** recommendations for estimating trends using smoothing splines.

| Consideration | Recommendation | mgcv nomenclature (if applicable) |
| --- | --- | --- |
| Capturing variation | 2-dimensional continuous-by-continuous smoother across year and day of year, implemented using a tensor product smoother | `te(doy, year, ...)` |
| Types of smoother | cubic regression smoother for year, cubic regression smoother or cyclic cubic regression smoother for DOY | `bs = c("cr", "cr")` or<br>`bs = ("cc", "cr")`<br>[used within `te()`] |
| How many knots? | One per year for year dimension, 5 for univoltine species, 7 for bivoltine species. With sufficient monitoring per year, more knots in the DOY dimension is generally better, but models will struggle or fail if there the fewest observations / year does not exceed the number of knots. | `k = c(5, length(unique(year)))`<br>[used within `te()`] |
| Special note for cyclic cubic regression smoothers | If using cyclic cubic regression smoother in the DOY dimension, you must specify the two DOYs that serve as endpoints for wrapping around. These are treated as two additional knots, so we recommend providing 5 knots for univoltine or 7 knots for bivoltine species, plus an additional 2 knots | `k = c(7, length(unique(year)))`<br>[used within `te()`]<br>`knots = list(doy = c(0.5, 365.5))`<br>[used outside of `te()`] |
| Anchor zeroes | In each year, add one artificial zero-count observation 20 days before the first observation across all years at the site and one 20 days after the final observation across all years at the site. |  |
| Calculating metrics | When calculating yearly metrics based on the activity curve, only use the activity curve predicted between the two anchor zeroes. |  |
| Estimating trends across years | When calculating growth rate, use all fitted years of data. Before calculating growth rate, set any estimated yearly abundance index below a small threshold (e.g. 0.5 or 0.1) to 0. It is then helpful to then add that threshold to ALL years to simplify fitting trends to log abundance index. When calculating phenology trends, exclude years with fewer than 2 non-zero counts. |  |
| Interpreting phenology trends of multivoltine species | Examine the activity curves with care, consider what estimated trends in phenology metrics actually represent. |  |

To support the use of our simulation framework, we have developed our code into an R package, ‘butterflyGamSims’ (https://github.com/cbedwards/butterflyGamSims), which can be used to (a) simulate and visualize butterfly monitoring data with known underlying population dynamics, and/or (b) analyze simulated or real data using a range of smoothing spline options. The package will be useful for ecologists seeking to identify the optimal modeling choices for a specific biological scenario or seeking to compare the efficacy of different monitoring schemes. Further, we hope the ability to simulate monitoring data with known properties will be useful for teaching or development of other methods. For details on simulation and analysis implementation, see the package documentation.

## Capturing year-to-year variation

In identifying population trends from monitoring data, there are two temporal dimensions to consider: variation between surveys *within* a year (hereafter, the “DOY” dimension, for Day Of Year) which defines the activity curve of that year and determines the yearly estimates of abundance and phenology, and variation *across* years (hereafter, the “year” dimension).

The simplest model for fitting monitoring data with a smoothing spline assumes that the shape and phenology of the activity curve (but not its height) is constant across years, and separately models a smoother in the DOY dimension and a linear trend in abundance across years (i.e., the global smoother scenario in Pederson et al. 2019). When fit using a log link, the estimated slope of the across-year trend is a measure of the population growth rate, accounting for overall seasonality. When modeling phenology as unchanging across years, this handily simplifies the estimation of growth rate. However, in many cases phenology is changing through time, especially in the context of climate change (Parmesan 2007). It is also possible that the shape of activity can change through time, in the form of changes in skew, the duration of activity, or even voltinism (Hodgson et al. 2011, Michielini et al. 2020). In any of these cases, it is more appropriate to model the activity curve in a way that allows it to vary between years.

When there are sufficient data within each year, fitting different activity curve smoothers for each year involves the least assumptions about relationships between years (Fig. 3B). This can be done with a continuous-by-factor smoother, which in mgcv is implemented with ‘s(DOY, by = ’year’)’. This allows the estimated activity curve for each year to be largely independent from one another, which will provide unbiased estimates of year-to-year variation. Unfortunately, because smoothing splines are highly flexible, a model can only fit each year independently if there is sufficient data in *every* year. In working with butterfly monitoring data in the United States, we have often encounter monitoring datasets with relatively few observations per year and individual years with substantially fewer days of monitoring. In these cases, we need a method that allows data-rich years to inform the estimated activity curve of data-poor years, so that we do not have to censor years of limited monitoring.

**Figure 3:**
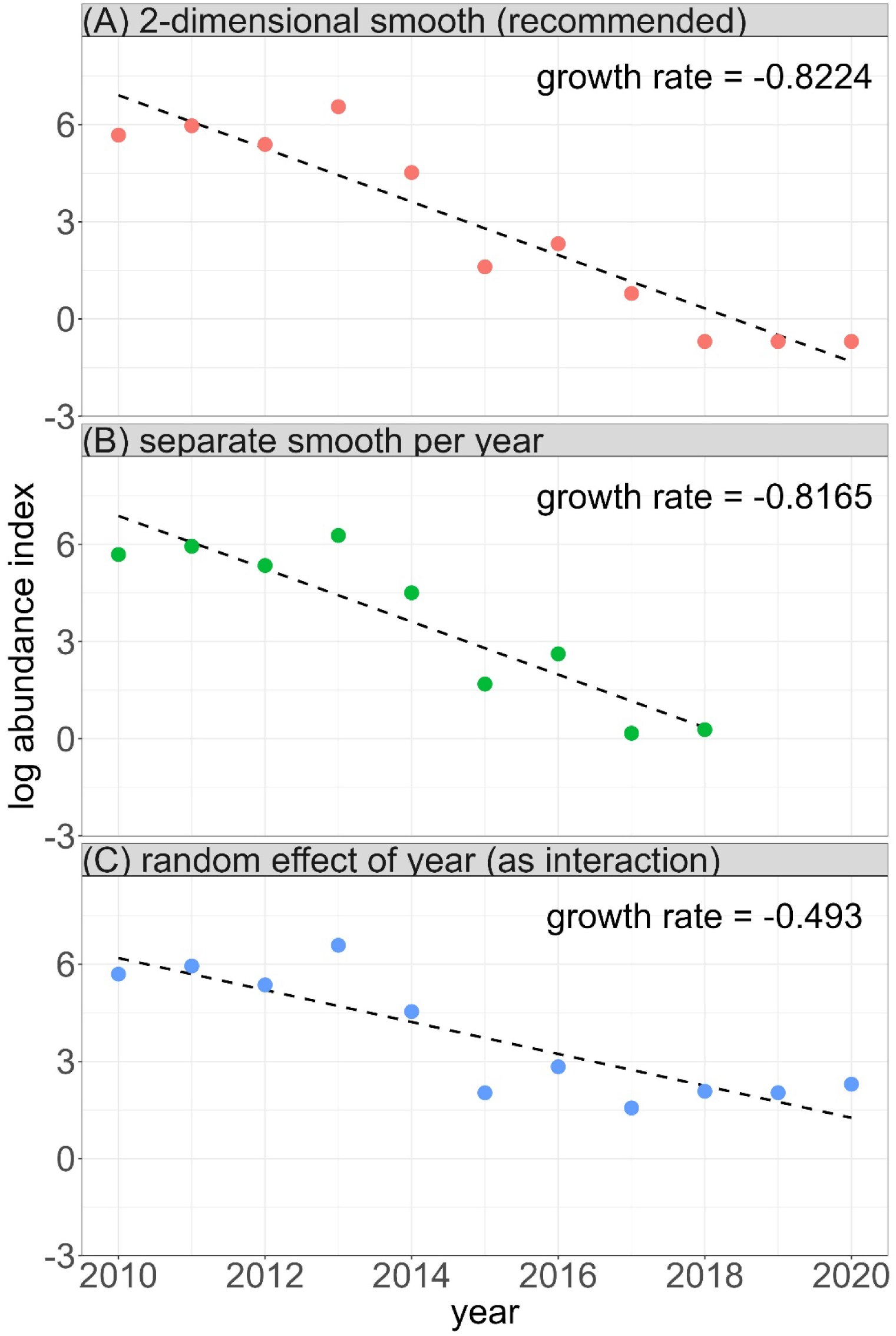
Three separate approaches for accounting for year-to-year variation in activity curve illustrated using Hermes Copper records at the “Sycuan Peak” site. A) The 2-dimensional smoother approach is our recommendation for estimating across-year trends in abundance and phenology, and allows nearby years to inform one another. This approach biases estimates towards reduced across-year variation, but should not bias estimates in trends. B) fitting each year with a separate activity curve involves the fewest assumptions, but the model failed to converge without censoring years with fewer than 4 records (2019 and 2020). C) Allowing the smoother to interact with a random effect of years allows data-rich years to inform data poor years, and does not bias years towards resembling adjacent years. However, the shrinkage associated with random effects pulls yearly estimates towards the mean, drastically underestimating declines.

Allowing information from each year to inform the fit of other years appears to be a problem well-suited to random effects, and it is possible to specify a continuous-by-random-effect smoother in a model (in mgcv, using ‘s(doy, year, bs = c(“cr”, “re”)’). However, using year as a random effect inherently biases estimates of across-year trends, as the shrinkage associated with the random effect will pull the years of highest and lowest abundance (and most extreme phenology) towards the across-year average. Consequently, using a continuous-by-random-effect smoother underestimates the magnitude of growths or declines (Fig. 3C). Instead, when seeking to estimate growth rates from sparse data, we recommend a 2-dimensional continuous-by-continuous smoothing term, with smooths across DOY and year. The interaction implicit in this smoother allows the activity curve to change in shape between years in such a way that data-poor years tend to resemble adjacent years (Fig. 3A). This approach biases results towards capturing less year-to-year variation, but so long as the assumption of linear trends across years is correct, this method does not produce biased estimates of trends. We note that when using a 2-dimensional continuous-by-continuous smoother across DOY and year, it is important to use a tensor product smoother (in mgcv, this is implemented with ‘te()’) instead of the default isotropic smoother (in mgcv, ‘s()’) (Pederson et al. 2019). A tensor product smoother allows smoothing across two or more variables with each variable having its own wiggliness penalty. As DOY and year are on wildly different scales, the appropriate smoothing across DOY is likely to be different from the appropriate smoothing across years.

## Specific implementation notes

Here we assume the reader is following our recommendation to apply a continuous x continuous tensor product smoother across DOY and year.

### What type of smoother?

Generalized additive models fit smoothing splines using a variety of basis functions, which can subtly or not so subtly constrain the shape and flexibility of the specified smoother. For two-dimensional smoothers, it is possible to specify different basis functions for each of the two dimensions, allowing us to treat the DOY and year dimensions separately. In the year dimension, we find a standard cubic regression basis performs well. In the DOY dimension, we can also use a cubic regression basis, but given the seasonal nature nature of DOY, a cyclic cubic regression basis may seem more appropriate (Pederson et al. 2019). A cyclic cubic regression spline is similar to a cubic regression spline, but constrains the beginning and end points of the fitted shape to match in value and slope, allowing us to explicitly specify our expectation that January 1 is one day away from December 31. In a naive model fitting (without our other recommendations), we found that using cyclic regression bases in the DOY dimension reduced the frequency of problematic model fits. However, using other appropriate methods as outlined below, we find that cubic regression and cyclic cubic regression splines perform similarly (Tables S3-5). Because cubic regression bases are slightly easier to work with (e.g., no need to specify endpoints), we recommend using them. We note that our simulations are for data in which the monitoring data does not span the entire year. In cases when data are available across the year such that the model can be expected to reasonably predict activity in early January and late December, we strongly recommend using a cyclic smoother (e.g., Canizares et al. 2023).

### How many knots?

Knots determine the maximum flexibility available to a fitted smoothing spline. Fitting with insufficient knots constrains fitted models to very simple shapes, while fitting with excess knots is more

computationally expensive and in practice can sometimes lead to overfitting (although the “smoothing” of smoothing splines tries to mitigate the risks of overfitting). When determining the appropriate number of knots in the year dimension, we found it was sufficient to use one knot per year. This allowed the fits of individual years to vary substantially if the data necessitated it, and when there was little year-to-year variability, the fitting process selected a stronger smoothing penalty, fitting a simpler shape across years. When determining the most appropriate number of knots across the DOY dimension, the best choice was less clear. Using butterflyGamSims, we found that there was generally little difference in performance based on knot number, but estimates for trends in the median for the Zonneveld activity curve (and likely other asymmetrical activity curves) were much more biased at the minimum 3 knots than higher values (Table S3). When estimating flight period for either univoltine activity curve, variance and error was lowest with 3 knots, but error and variance were not substantial even with 5 knots. Since we generally don’t expect to know the exact shape of activity curves, we recommend using 5 knots for univoltine species where possible. For simulations of bivoltine activity curves, for our two baseline scenarios, we saw a small improvement in mean error for all estimated metrics when using 7 knots rather than any other number between 6 and 10. However, this benefit disappeared when modify the growth rate (Table S5). Nonetheless, without any clear best option, 7 knots is a reasonable choice for bivoltine species. If readers suspect a particular biological scenario or shape of activity curve for their data, they can simulate data with those properties and compare bias and variability for their specific use case with butterflyGamSim.

### Avoiding misfits

When working with sparse data, model fits can sometimes produce biologically unreasonable results (e.g., Fig. 5A, year 2012). As an extreme example, if we were fitting monitoring data from a single year, if monitoring ended too early relative to butterfly activity and we have only data before and up to the peak of activity, the smoothing spline could predict continually increasing activity on to the end of the year. Using a 2-dimensional smoother can sometimes prevent these kinds of unreasonable fits, as the model will favor a shape that is similar to adjacent years. So long as most years have monitoring data throughout the activity period, this can prevent unreasonable fits in the remaining years. However, issues can still arise, especially in the initial or final years of a data set, which have fewer adjacent years of data to inform the fit. In addition to choosing appropriate smoother types and knots counts, we found two methods to be effective at minimizing the frequency and consequences of these catastrophic misfits: adding anchor zeroes and limiting the period of calculation. Fundamental to these approaches is the understanding that there are constraints on the timing of butterfly activity for temperate species. For example, the Fender’s Blue Butterfly is active between mid April and early July (Bonoan et al. 2021), and there is no reasonable possibility of adult activity in December and January.

To capture our biological intuition for the limits of activity, we follow Rothery and Roy (2001) in adding artificial “anchor zeroes” before and after the actual time series data each year. These new zero count observations favor model fits in which predicted activity falls to zero before and after the actual monitored activity period. Anchor zeroes can be added at any DOY for which we are certain monitoring would report 0 activity, but doing so too far from the actual monitoring data reduces the efficacy of this approach. Placement of anchor zeroes can be based on expert opinions or independent studies, but a simple implementation is to place them a fixed distance before and after the earliest and latest DOY of monitoring of a given population. Monitoring is generally informed by on-the-ground observations of the population, and using the range of monitored days as a baseline leverages the expertise and knowledge of the field biologists carrying out the monitoring. The **rbms** R package, which was developed for fitting data from the UK Butterfly Monitoring scheme, defaults to adding seven anchor zeroes on either side of the monitoring season, placed daily from one to seven days before and after the season starts and ends (Schmucki et al. 2022). However, for working with sparse, less consistent monitoring data, we recommend adding a single anchor zero rather than seven, with a several-day buffer to account for the possibility that monitoring missed part of the activity. Our simulations showed generally somewhat better performance using anchors zeroes placed 20 days before and after the final days of monitoring (rather than 10 days). Using anchor zeroes at either 10 or 20 days considerably reduced variation between simulation estimates for Zonneveld and bivoltine activity curves relative to not using anchor zeroes (Fig. 4A). We note that our simulation findings are very conservative, as we simulated data in which monitoring spanned the full activity curve every year; in practice, anchor zeroes are most valuable when fitting data in which data in some years have only partial coverage of the activity curve.

**Figure 4:**
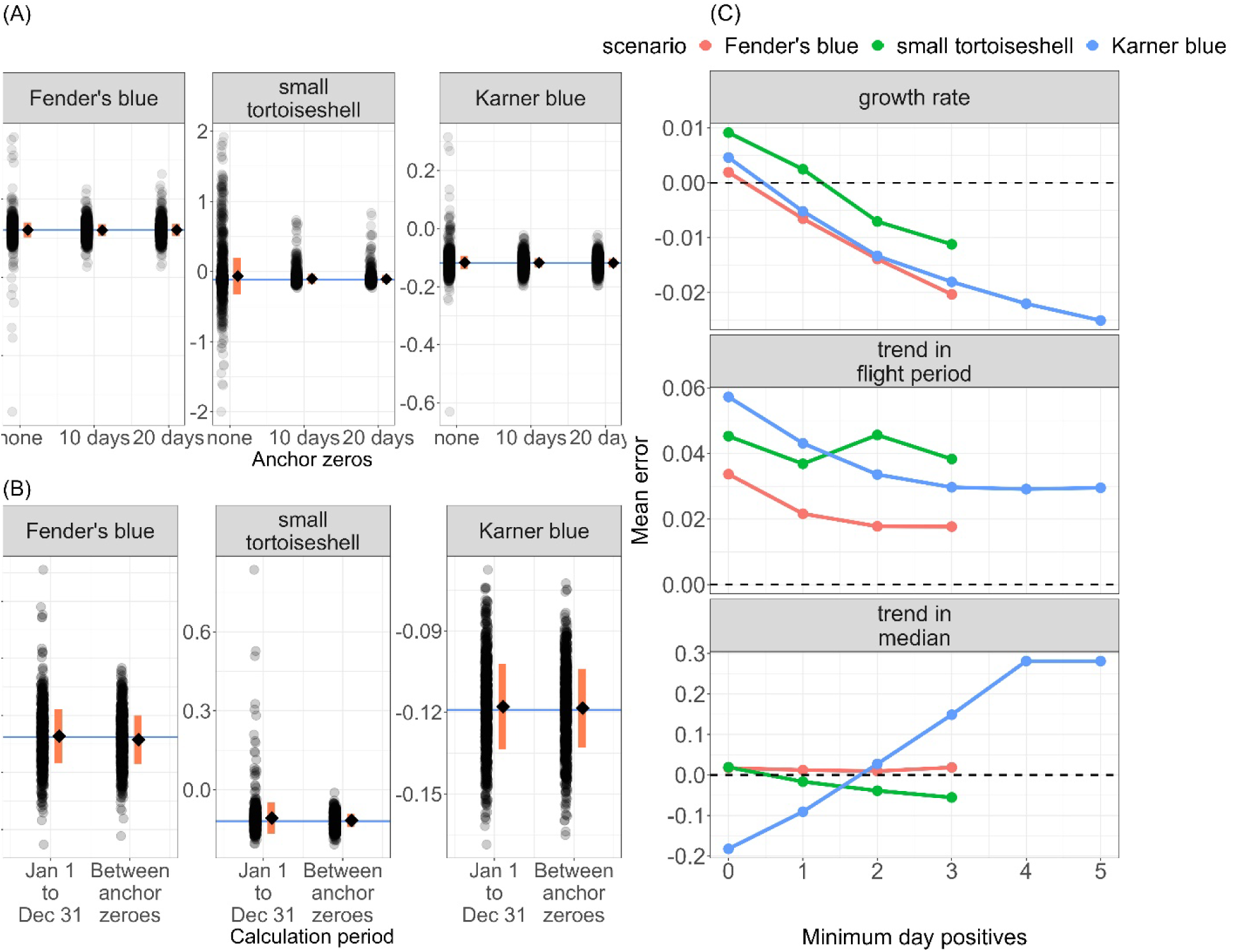
Results of ButterflyGamSim simulations parameterized for three butterfly species show the importance of several methods. (A) Adding anchor zeroes reduces the variability between simulations, and prevents rare instances of extremely wrong estimates. Transparent black points: 1000 individual simulation results for each parameter combination; diamonds: average simulation results; orange bar: +/- 1 standard deviation; horizontal line: true growth rate. (B) Limiting the calculation of activity curves and yearly metrics to the period between anchor zeroes reduces the impacts of rare instances of extremely poor yearly fits. (C) Excluding years with 0 or 1 day positives can reduce bias in estimating phenological trends, but generally increases bias when estimating growth rates. Points show the average across 1000 simulations, excepting simulations which had insufficient years of data to estimate trends after applying the exclusion rule; dashed line shows the goal of a mean error of 0 (no bias).

The use of anchor zeroes greatly reduces the frequency of catastrophic misfits (Fig. 5B). A complementary approach is to minimize the harm of misfits by only calculating metrics based on predictions of the fitted model within a limited and biologically reasonable range. In doing so, poor model behavior outside the biologically relevant period will not affect estimated population metrics. For simplicity, we recommend using the period defined by the anchor zeroes, as those are already chosen with the goal of spanning any biologically reasonable activity. We found that doing so substantially reduced variance between simulation estimates (e.g., Fig. 4B), especially when estimating trends in phenology (Table S2).

**Figure 5:**
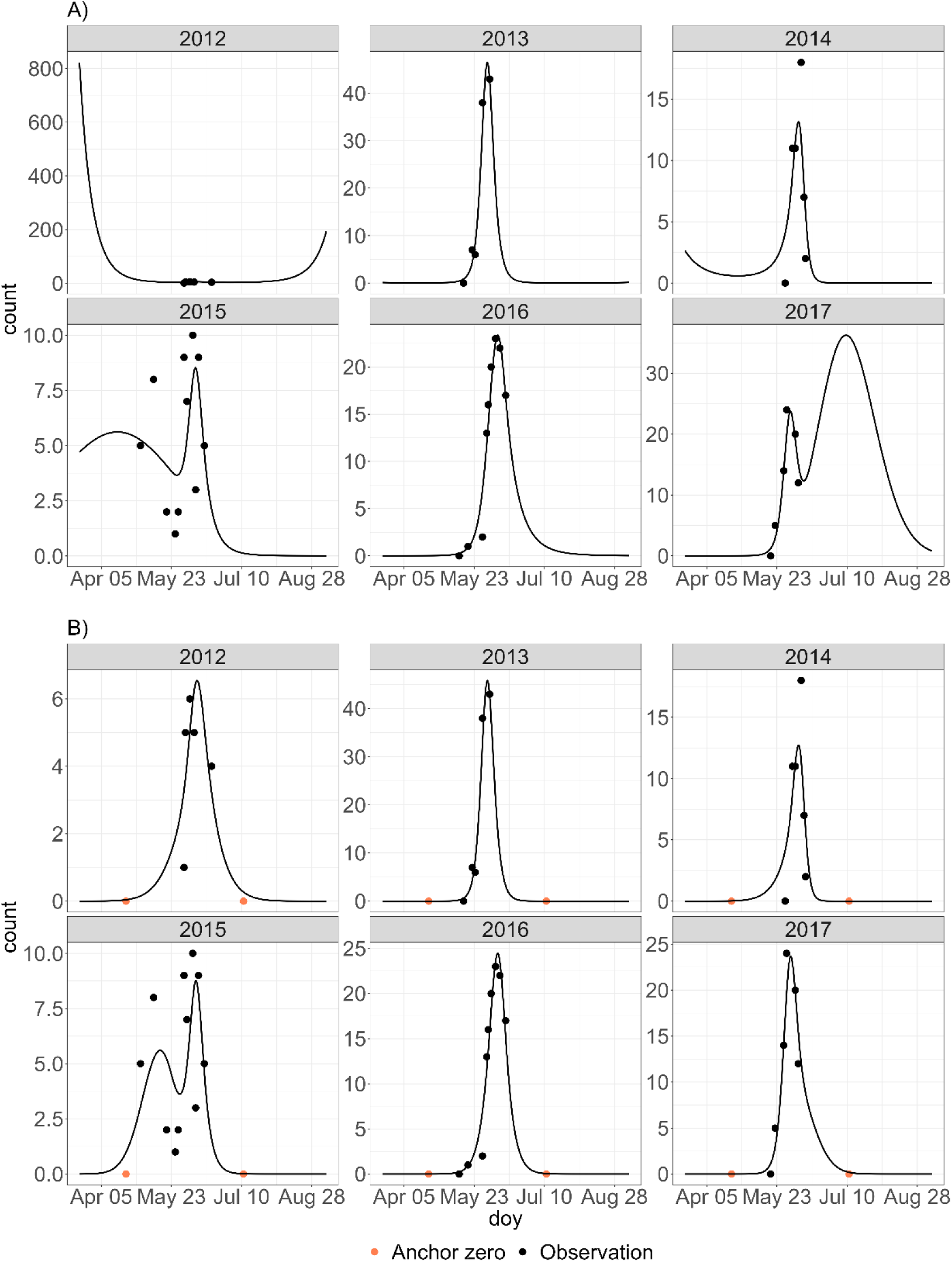
Smoothing splines sometimes produce “catastrophic misfits” that qualitatively misrepresent butterfly activity. The use of anchor zeroes can prevent this. (A) Observations of Hermes Copper at Boulder Creek site, and smoothing spline fit to all years of data (years 2012-2020), with the model structured based on our recommendations but not including anchor zeroes. 2012, 2014, 2015, and 2017 exhibit catastrophic misfits for which estimated phenology and abundance will be unreasonable. (B) By adding “anchor zeroes” – dummy observations with 0 count (coral points)– 20 days before and 20 days after the earliest and latest real observations in the data set, we represent our expectation that field biologists time their monitoring to the period when butterflies are active. Refitting this data set with anchor zeroes avoids each of the observed catastrophic misfits.

Even with careful choice of model parameters and the use of anchor zeroes, we find that especially messy and sparse data can still occasionally lead to catastrophic misfits. When fitting monitoring data with smoothing splines, it is important for ecologists to examine the predicted activity curve for each year of data and exclude any years with biologically impossible results that produce meaningfully incorrect metrics from post-hoc analysis.

## Estimating trends across years

Once the smoothing spline is fit and individual yearly metrics are calculated, we can still encounter unreasonable results when individual years -- or periods of years -- have few or no non-zero counts (“day positives”, sensu Casner et al. 2014). Whether the butterfly was truly absent or merely rare enough that monitoring failed to observe it, these years caused problems. When years with no (or few) day positives are part of a larger time series fit with 2-dimensional smoothers, the activity curves for those individual years will tend to be extremely flattened out, with very early estimated onset, very late estimated ends of activity, flight periods that are subsequently very long, and median days of activity that are drawn towards the middle of the predicted period (i.e., the median day for a completely flat activity curve). Including these all- or mostly-zero-count years can lead to unreasonable estimates of phenology trends for populations that either become extirpated or recover from near extirpations. In our work, for several populations that appeared to become extirpated, we initially estimated their flight periods were increasing by approximately 100 days per decade. Clearly it is important to exclude these problem years with wildly unreasonable phenology metrics when estimating phenology trends. On the other hand, it is important to include every year that is not biasing estimated trends, both to increase the data available for fitting and because years of low abundance may exhibit very real differences in phenology. Using simulations of populations trending towards extinction, we found that bias was generally minimized by excluding years of fewer than 2 non-zero counts when estimating trends in phenology (Fig. 4C). In contrast, *all* years should be included when estimating trends in abundance, as excluding years consisting mostly or entirely of zero counts inherently biased estimates of trends in abundance.

Despite the importance of including years of only zero counts when estimating abundance, a non-obvious problem arises for populations that end (or start) with several years of all zero observations. Smoothing across years favors model fits in which adjacent years are more like each other than the data of each individual year would otherwise suggest. Consequently, when a population declines and ends in several years of only zero observations, we frequently encountered cases where each of the years of all-zeroes had tiny but non-zero estimated abundance indices (e.g., less than 0.1 observable individual-days), with the later all-zero years having proportionally much *smaller* estimated abundance indices than the earlier ones (Fig. 6). In practical terms, a fitted model that estimates abundances indices of 0.1, 0.01 and 0.001 for the final three years of a population time series is predicting that there is no reasonable expectation of observing individuals in any of those years using the current monitoring scheme. This implies the three years are largely equivalent in underlying biology. However, fitting a linear model to log abundances for those three years will lead to a very strong signal of population decline with an estimated growth rate of −2.3 (90% decline per year). In other words, *after the population appeared to be extirpated or undetectable*, *we estimated substantial declines*. Further, the tendency for estimated years to resemble adjacent years (the property of gams which allows non-zero abundance earlier in a time series to “leak” into the years of all-zeroes) is not consistent from one dataset to another and depends on the model smoothing parameter that are estimated from the data. Consequently, estimated growth rates for populations that appear to become extirpated can vary substantially based on the underpinnings of GAM algorithms and the variability (or lack thereof) between earlier years in the monitoring program. These sources of variation do not reflect biological processes and are especially problematic when using non-parametric bootstrapping to estimate uncertainty. In our work on populations with apparent extirpations, bootstrapped abundance indices generally resembled the estimates of the real data for years containing at least one non-zero survey but varied widely in how small the “very small” estimates of abundance were for years of only zero counts (Fig. 6).

**Figure 6:**
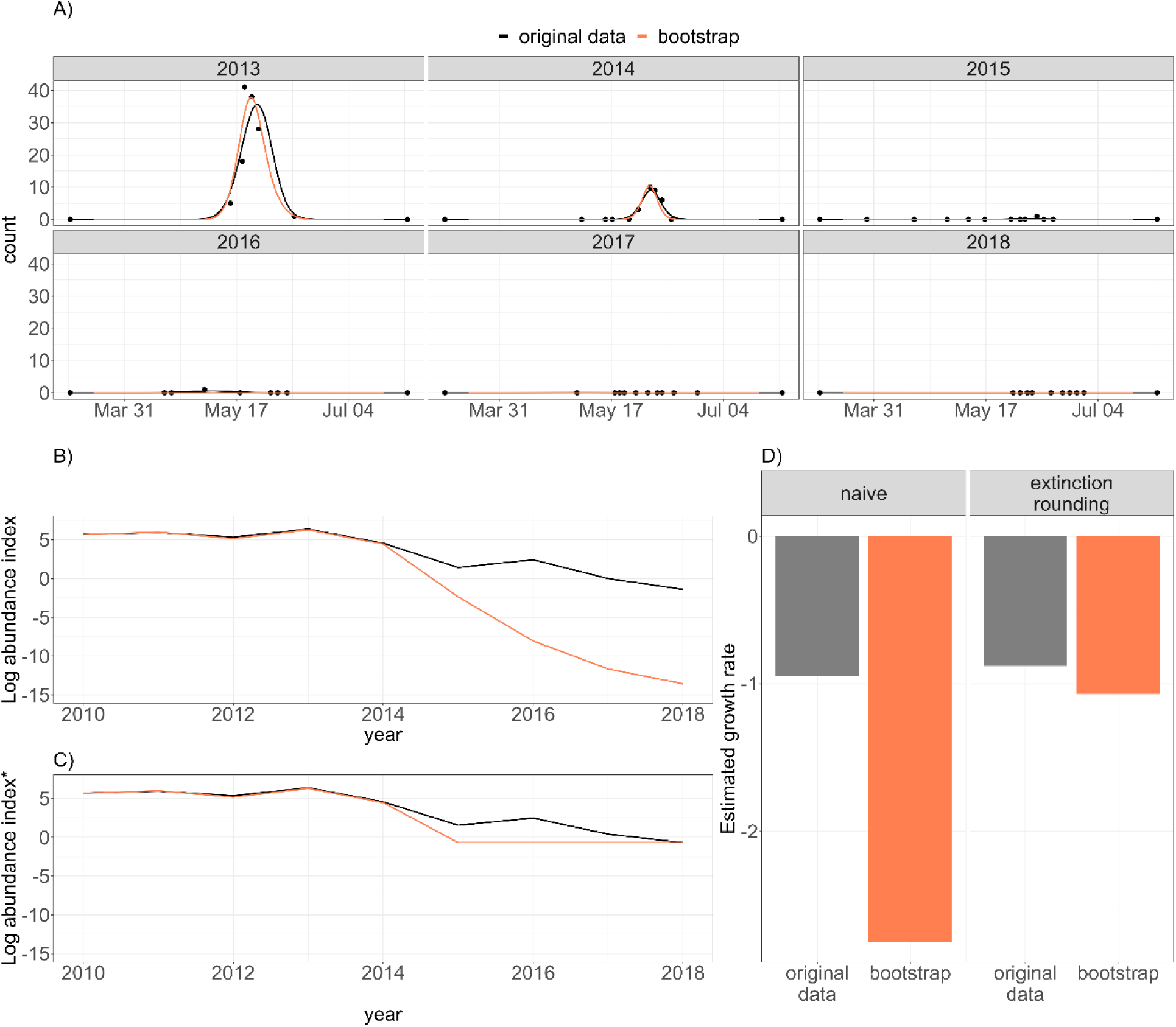
When populations fall to apparent extinction, miniscule differences in model fit can lead to extreme differences in estimated growth rate. Here we illustrate this with a model fit to the Hermes copper population at Sycuan peak, and model fit to bootstrapped data which resampled within years, which should produce similar yearly metrics. (A) Data (black points) and fitted models (curves). Black lines show predictions of model fit to all the data; coral lines show predictions from a bootstrap replication. The two fitted models show qualitatively similar patterns, predicting high activity in 2013, declining to effectively no activity in 2016 and after. (B) However, on a log scale, abundance indices differ substantially between the original fit and the bootstrap for the years in which data was entirely 0s (2016 and onward). (C) By setting all abundance indices below some small threshold value (here 0.5) to the threshold value, years that qualitatively represent extinction are treated as equivalent. This reconciles timeseries that diverge on a log scale in years of very low abundance. (D) Without this rule, the rate of decline for the bootstrap timeseries was more than twice that of the original fit, despite having nearly identical abundance indices before 2016. When applying our thresholding rule, the two models produce similar estimated growth rates.

We recommend a solution borrowed from theoretical ecology, in which many population models are mathematically constrained such that abundance can approach 0, but never reach it. In those cases, theoretical ecologists often identify a minimum threshold for non-zero values and treat anything below that as zero. Similarly, when estimating growth rate from yearly estimates of abundance index, we recommend rounding any sufficiently small abundance index (e.g., below 0.5 or below 0.1) to 0 (Fig. 6D). In effect, we are adding a rule to define populations as “apparently extirpated” in individual years with sufficiently small estimated abundance indices, and then treating all of these years as having an equal abundance (of 0). For fitting purposes, this then necessitates adding a small constant to *all* abundance indices to fit a log-linear trend, as is common practice for fitting the log of data containing zeros.

### [BOX:] Example analysis protocol

1. Start with a single population of monitoring data, with multiple survey days per year, and sufficient years to estimate across-year trends of interest.
2. Augment these data with anchor zeroes. For each year, add one additional artificial observation of zero individuals 20 days before the earliest day of monitoring in all years, and another artificial observation of zero individuals 20 days after the last day of monitoring in all years. (from “avoiding misfits”)
3. Fit the data using a smoothing spline with a tensor product smoother across DOY and year (with year as a continuous variable not a factor), using a cubic regression basis in each dimension. In the year dimension, use one knot per year; in the DOY dimension use 5 knots (for univoltine species) or 7 knots (for bivoltine species). If data for a univoltine species is so sparse that this model will not fit, use 3 knots in the DOY dimension instead (and recognize that estimated trends in median phenology may be somewhat less reliable) (from “Specific implementation notes”).
4. For each year of data, use the fitted model to predict the count on an evenly spaced sequence (e.g., every 0.1 days) from the day of the initial anchor zero to the day of the final anchor zero. Using these predictions (which are the estimated activity curve), calculate the abundance index for each year as the sum of predicted counts multiplied by the number of days between each prediction (e.g., 0.1). Calculate any phenology metrics of interest using the cumulative sum of the predicted counts counts to easily find the quantiles. (from “avoiding misfits”)
5. To estimate population growth rates, round any abundance index below 0.1 to 0. Add 0.1 to *every* estimated abundance index to make it possible to take the log of abundance indices. Fit a linear model to the log of abundance indices. (from “estimating trends across years”)
6. To estimate trends in phenology across years, fit a linear model in which each year’s estimate of the phenology metric is a separate data point. Only include years in which the original data had 2 or more day positives. (from “estimating trends across years”)
7. To estimate uncertainty for any across-year trends of interest, use non-parametric bootstrapping of the original data (resampling within each year), repeating steps 2-6 for each bootstrapped instance. This provides bootstrapped distributions for estimated growth rates and trends in phenology, and bootstrapped iterations can be used to characterize uncertainty of any further analyses.

### Cautions

We include a few warnings about using smoothing splines to infer trends in abundance and phenology from monitoring data, motivated by our own experiences.

First, with more complex, hierarchical data, it can feel intuitive to integrate data from multiple populations into a single model. For example, with multiple sites for a single species, we could fit a model with a smoother across day of year separately for each site (in mgcv, using ‘by = sitè), with a separate smoother of DOY by year to capture our expectation that sites vary but likely have some similarity in activity in each year (e.g., there are good years and bad years for butterflies). In our experience, adding these complexities can increase the frequency of model misfits. Further, these additional terms can lead to constraints in the model fit that do not reflect biological reality. For example, depending on the spatial scale between sites and the drivers of population dynamics, populations may be largely independent and there may *not* be good years and bad years for the joint data as a whole. Similarly, adding spatial dimensions to smoothers is technically possible, and doing so is tempting as it enables the model to estimate continuous geographic variation in abundance and growth rates. However, when developing and exploring DOY by latitude by longitude smoothers for hundreds of butterfly species across North America, we discovered that smoothing across space prevented models from capturing any meaningful variation in growth rates across space, and the predicted activity curves were frequently biologically impossible (e.g. inverted activity curves) at the edges of modeled spaces and in regions of low data density.

Second, trends of standard phenology metrics can be difficult to interpret in the context of multivoltine activity curves, particularly when individual generations are distinctly separated as is common for some bivoltine butterfly species (Fig. 7). As Inouye et al. (2019) highlights, the population activity curve should still be interpreted as the activity of many individuals, and numerous biological processes on the scale of individuals can explain an advance in population-level phenology. However, when we identify an advancing median phenology for a univoltine species, we can report with some reasonable confidence that, in general, individuals are active sooner. In contrast, an advancing trend in median phenology from a bivoltine species can be caused by (a) both generations shifting the same amount, (b) one generation advancing while the other remains static (or even retreats to a lesser degree), (c) both generations remaining static in phenology, but the initial generation increasing in abundance relative to the second generation, or (d) some combination thereof (Fig. 7B-D). We have encountered cases like this in our analyses, in which the relative abundance of the first and second generations change and the estimated median consequently jumps tens of days in the span of a year (e.g. Figure 7A). Similar issues can arise with estimates of onset, end of season, and flight period if the relative abundance between generations shifts to the point that the 0.1 and 0.9 quantiles become associated with a single generation (or stop being associated with a single generation).

**Figure 7:**
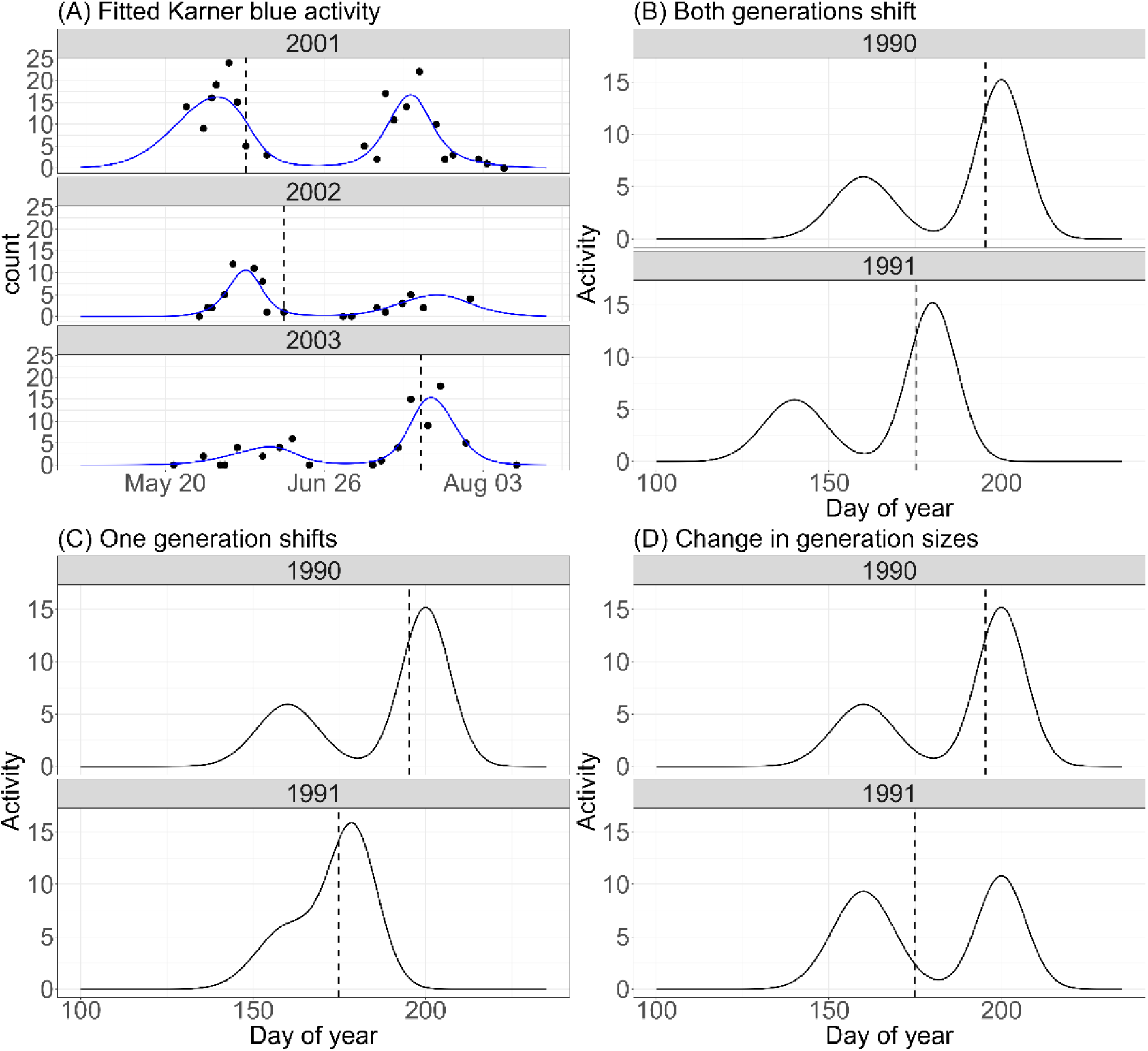
For bivoltine species, differences in phenology metrics between years can reflect multiple processes. (A) fitted data for Karner Blue at a site in the Albany Pine Bush Preserve: black points show counts of observed butterflies, blue lines show predicted activity curve, dashed line shows median activity. Estimated median activity changes by more than 40 days in three years, even though neither generation shifts that much. (B-C) Different biological processes can produce identical phenological shifts. Here, an advance in median phenology of 20 days is caused by (A) both generations advancing by 20 days, (B) one generation remaining constant, the second advancing by 21 days, and (C) each generation maintaining the same phenology, but shifting in relative abundance. Interpreting phenological trends for bivoltine species requires caution and careful examination.

We do not have a general solution to the challenges of interpreting phenology metrics for multivoltine species, and we hope to see further research and development in this area. For specific cases, it may be reasonable to use or develop more appropriate metrics that are less ambiguous. For example, analyses of distinctly bivoltine butterfly species could calculate the standard metrics we propose separately for each generation (e.g., separate median phenology and associated trends for each generation). This likely would not work for species with overlapping generations, however. Further, a strength of the metrics we emphasize (median, onset, end of activity, flight period) is that they *can* be calculated and compared for a wide range of activity curves, making it possible to examine phenology trends across species without needing to create incompatible separate metrics for species with different life histories. For now, we merely caution readers to think carefully -- and examine their fitted models -- before relying too heavily on the estimated phenology trends of multivoltine species.

## Conclusion

Broad-scale insect declines of the Anthropocene have highlighted the importance of estimating species trends, and ongoing interest and research into population responses to climate change frequently necessitate estimating phenological trends (Dirzo et al. 2014, Goulson 2019, Van Klink et al. 2020). Ecologists are harnessing new and previously overlooked data sources to better understand populations in our changing world (e.g., Gotelli et al. 2021, Ellwood et al. 2022, Di Cecco et al. 2023), and smoothing splines have proven a useful tool in inferring population and phenology trends from insect monitoring data (Hodgson et al. 2011, Wepprich et al. 2019, Stemkovski et al. 2020). However, rare and at-risk species are frequently under-represented in comparative studies of insect trends, due in part to the limited and messy data available for such species and the data requirements of standard methods (Forister et al. 2023). We share our recommendations in the hopes that they will facilitate analyses of previously unusable data, and better inform our understanding of insect population dynamics and phenology. Further, we hope that the ButterflyGamSims package proves useful to ecologists interested in testing or adapting their own methods, educators looking to generate reasonable datasets with known properties, and habitat managers looking to explore the efficacy of different monitoring regimes.

## Data Accessibility Statement

Scripts, simulated data and the butterfly data that feature in the examples will be made available on Figshare. The butterflyGamSims package is available at https://github.com/cbedwards/butterflyGamSims.

## Author contributions

**Collin B. Edwards**: Conceptualization (lead); data curation (lead); formal analysis (lead); software (lead); visualizations (lead); writing – original draft (lead); writing – review and editing (equal)

**Cheryl Schultz**: data curation (support); visualizations (support); writing – review and editing (equal) **David Sinclair**: formal analysis (support); visualizations (support); writing – review and editing (equal) **Noam Ross**: formal analysis (support); visualizations (support); writing – review and editing (equal) **Dan Marschalek:** Investigation (lead); writing – review and editing (equal)

**Elizabeth Crone**: formal analysis (support); visualizations (support); writing – review and editing (equal)

## Acknowledgements

This work was conducted with support from the DOD SERDP program (RC-2700), and partial support for

C.B.E. was provided by the USGS John Wesley Powell Center for Analysis and Synthesis Working Group on Status of Butterflies (G22A002192) and the USFWS. We thank Steve Campbell for the use of their Karner blue data in our examples and figures. We thank Jani Uusitalo and Aaron Carlson for their photos of Fender’s blue and small tortoiseshell butterflies.

## Supplements: Simulation results

We simulated 1000 iterations each of 1584 combinations of parameter values in a factorial design (3 anchor options x 2 limitation options x 2 smoother types x 3 or 5 knot counts x 6 exclusion criterion x 3 butterfly scenarios x 2 rates of decline), for which we estimated trends in multiple metrics (growth rate, median phenology, onset, flight period); we then carried out additional targeted simulations to illustrate individual points and test for sensitivity to population growth rates. Presenting every aspect of these results would be unreadable. We instead report how individual decisions influence model fit when other parameter values are at optimal or reasonable baselines (with the exception of testing the effect of anchor zeroes, explained below). Modeling choices can improve model fit in two ways. First, good choices sometimes make estimates from different simulations of the same parameters more similar to one another; this decreased variance, and in application means estimates from the model-fit will be more reliable. Second, good choices sometimes bring the average estimates across simulations closer to the true trends; this decreased bias, and in application means that we don’t expect our estimates to be consistently incorrect in the same direction. Below we report the average error in estimates (a measure of bias) and the standard deviation of estimates (a measure of variance), in all cases using 1000 simulations for each combination of parameter values.

### Anchor zeroes

Adding anchor zeroes substantially improved model fit in many cases, decreasing the standard deviation of simulation estimates by as much as 4-fold. Using anchor zeroes at 20 days was generally more effective than at 10 days, especially for more extreme growth rates, but the difference between 10 and 20 days was much smaller than between using and not using anchor zeroes. Note that because we were comparing simulations with anchor zeroes and without, it was not reasonable to limit the calculation of metrics to the period defined by the anchor zeroes. In all cases the calculation of metrics is based on predictions from day 1 to day 365.

**Table S1:** Average error (bias) and standard deviation of estimated trends (variance) for different anchor choices.

| Activity curve | Anchors | Growth rate |  | Trend in median mean |  | Trend in flight period |  |
| --- | --- | --- | --- | --- | --- | --- | --- |
|  |  | mean error | sd | error | sd | mean error | sd |
| Fender's blue (Gaussian) | None | 0.0007 | 0.028 | -0.0038 | 0.361 | 0.0288 | 0.099 |
|  |  |  | 6 |  | 0 |  | 3 |
|  | 10 days | 0.0000 | 0.022 | 0.0025 | 0.211 | 0.0204 | 0.083 |
|  |  |  | 5 |  | 2 |  | 3 |
|  | 20 days | 0.0007 | 0.023 | 0.0169 | 0.304 | 0.0300 | 0.110 |
|  |  |  | 8 |  | 0 |  | 3 |
| small tortoiseshell (Zonneveld) | none | 0.0527 | 0.260 | 0.3887 | 2.224 | 0.1943 | 0.285 |
|  |  |  | 1 |  | 9 |  | 1 |
|  | 10 days | 0.0156 | 0.069 | 0.2445 | 0.947 | 0.0743 | 0.177 |
|  |  |  | 2 |  | 1 |  | 5 |
|  | 20 days | 0.0118 | 0.060 | 0.1789 | 0.767 | 0.0759 | 0.176 |
|  |  |  | 2 |  | 9 |  | 5 |
| Karner blue (Bivoltine) | none | 0.0024 | 0.023 | -0.1083 | 0.636 | 0.0418 | 0.124 |
|  |  |  | 4 |  | 9 |  | 8 |
|  | 10 days | 0.0011 | 0.016 | -0.0493 | 0.483 | 0.0275 | 0.094 |
|  |  |  | 0 |  | 0 |  | 3 |
|  | 20 days | 0.0011 | 0.015 | -0.0393 | 0.449 | 0.0297 | 0.095 |
|  |  |  | 7 |  | 1 |  | 2 |

*Other parameters: Growth rate = -0.12, cubic regression basis in DOY dimension; 5 knots for univoltine species, 7 for bivoltine species, not limiting to anchor, 5 days of monitoring for univoltine species, 10 for bivoltine species; no excluding individual years based on day-positives.*

### Limiting calculation to anchor zeroes

Calculating metrics based on activity curve predictions from the first through the second anchor zero was consistently better than calculating based on predictions for the entire year. When estimating growth rates, we find a general reduction in the variance between simulations when applying this limit; further, we no longer observe extreme outliers representing individual simulations with one or more wildly implausible yearly estimates (Fig. 4B). Because the asymmetrical Zonneveld activity curves had skewed variation in growth rate estimates without the limit (unusually poor fits generally estimated overly positive growth rates), using the limit also reduced bias. When estimating phenology metrics, we consistently saw improved performance in terms of variance between simulations, typically quite large (e.g., an almost 10-fold reduction in the standard deviation of estimated trends in flight periods for Gaussian activity curves).

**Table S2:** Average error (bias) and standard deviation of estimated trends (variance) when either limiting the activity curve calculation to the period between anchor zeroes, or calculating the activity curve for the entire year.

| Activity curve | Limit? | Growth rate |  | Trend in median |  | Trend in flight period |  |
| --- | --- | --- | --- | --- | --- | --- | --- |
|  |  | mean error | sd | mean error | sd | mean error | sd |
| Fender's blue (Gaussian) | No |  | 0.023 |  | 0.304 |  | 0.110 |
|  |  | 0.0007 | 8 | 0.0169 | 0 | 0.0300 | 3 |
|  | Yes |  | 0.021 |  | 0.102 |  | 0.014 |
|  |  | -0.0023 | 1 | -0.0035 | 1 | 0.0087 | 2 |
| small tortoiseshell (Zonneveld) | No |  | 0.060 |  | 0.767 |  | 0.176 |
|  |  | 0.0118 | 2 | 0.1789 | 9 | 0.0759 | 5 |
|  | Yes |  | 0.024 |  | 0.093 |  | 0.033 |
|  |  | 0.0023 | 8 | -0.0099 | 6 | 0.0159 | 1 |
| Karner blue (Bivoltine) | No |  | 0.015 |  | 0.449 |  | 0.095 |
|  |  | 0.0011 | 7 | -0.0393 | 1 | 0.0297 | 2 |
|  | Yes |  | 0.014 |  | 0.419 |  | 0.021 |
|  |  | 0.0006 | 5 | -0.0209 | 6 | 0.0134 | 9 |

Other parameters: As in table S1, but using anchor zeroes at 20 days.

### Knots and smoothers

The choice of smoother did not substantially affect the average error or variation among estimates of growth rates. With appropriate anchors (20 days) and period for calculation (limit.to.anchor = TRUE), we saw no consistent advantage between using cyclic cubic regression smoothers and cubic regression smoothers across the simulated number of knots (Table S3, S4).

In general, the number of knots had limited effects on estimated growth rates. Depending on the population dynamics (Table S3 vs S4 vs S5) and the activity curve, mean error and the standard deviation differed slightly based on the number of knots, but bias was always less than 0.01 and variance remained fairly consistent for knots and smoother types with the same population dynamic and activity curve. For the Fender’s blue simulations (Gaussian activity), error in phenology trends was also minuscule in all cases, with relatively lower error and variance for lower knot counts. In part, this likely reflects the fact that a 3-knot gam can exactly recreate a gaussian curve, while gams with more knots can still recreate a gam but may also find other shapes to be more plausible for individual data sets. For the small tortoiseshell simulations (Zonneveld activity), we see substantial bias for trends in median phenology when using 3 knots (the average trend was off by more than 1 day / decade); this bias fell with increasing knots, and was generally an order of magnitude lower at 5 knots. For the Karner Blue (bivoltine activity), average error in median trends and flight period were relatively low, but both were generally at their lowest with 7 knots. This may be an artifact of the specific way in which the bivoltine distribution is specified (it is composed of two gaussian curves, and 3-knot gams can exactly recreate a gaussian curve). However, this generally supports 7 knots as a choice for bivoltine activity curves.

**Table S3:**
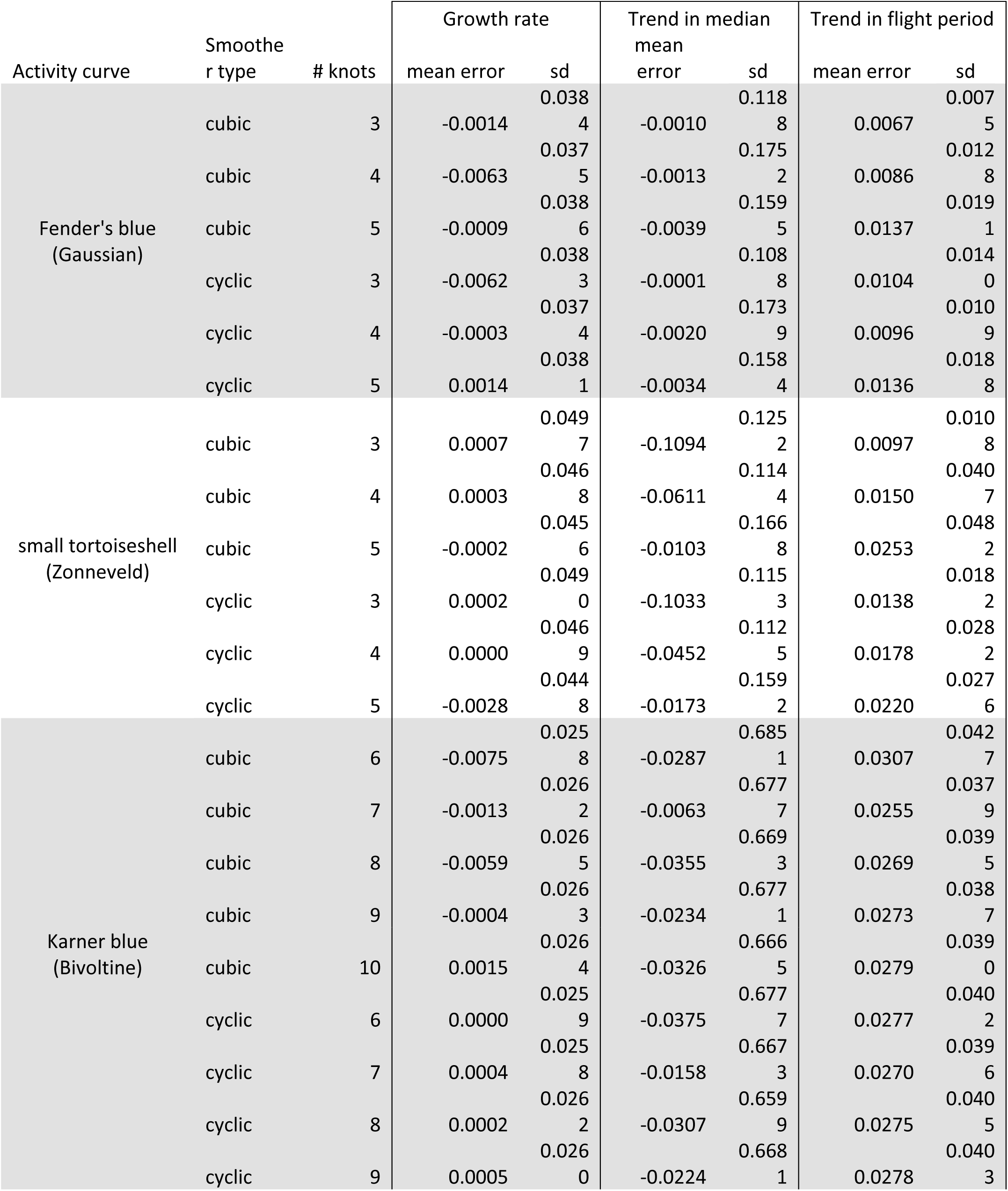
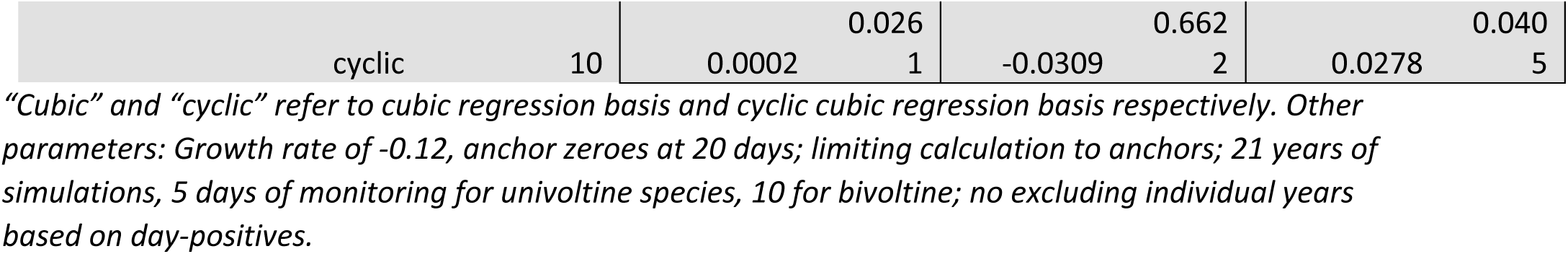
Average error (bias) and standard deviation of estimated trends (variance) when fitting models using one of two smoother types and one of a range of knot numbers in the DOY.

**Table S4:**
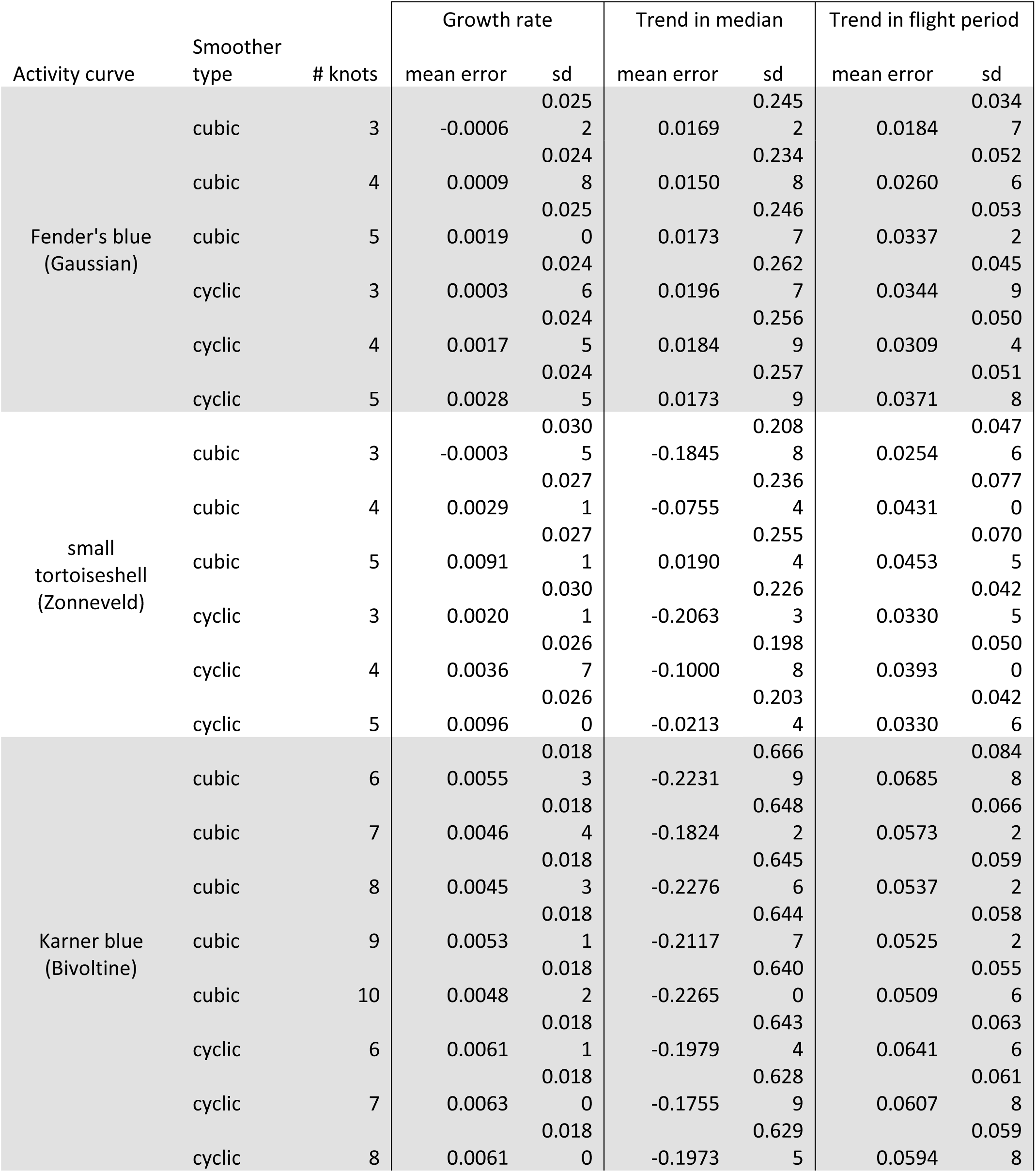
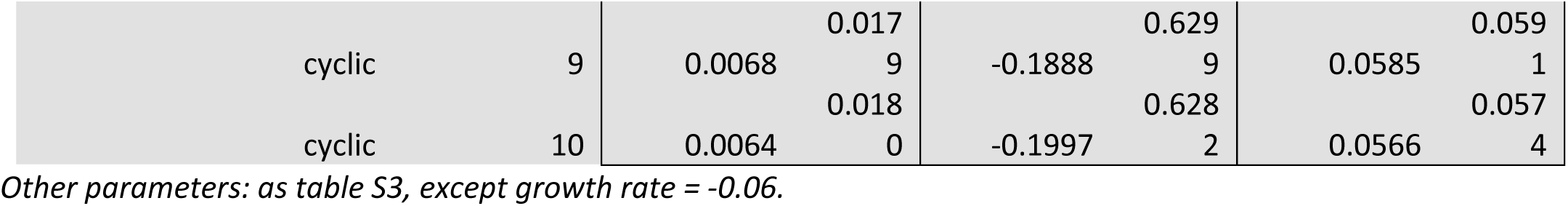
Average error (bias) and standard deviation of estimated trends (variance) as in Table S3, but using more extreme declines and longer duration, designed to emulate populations reaching extirpation.

**Table S5:**
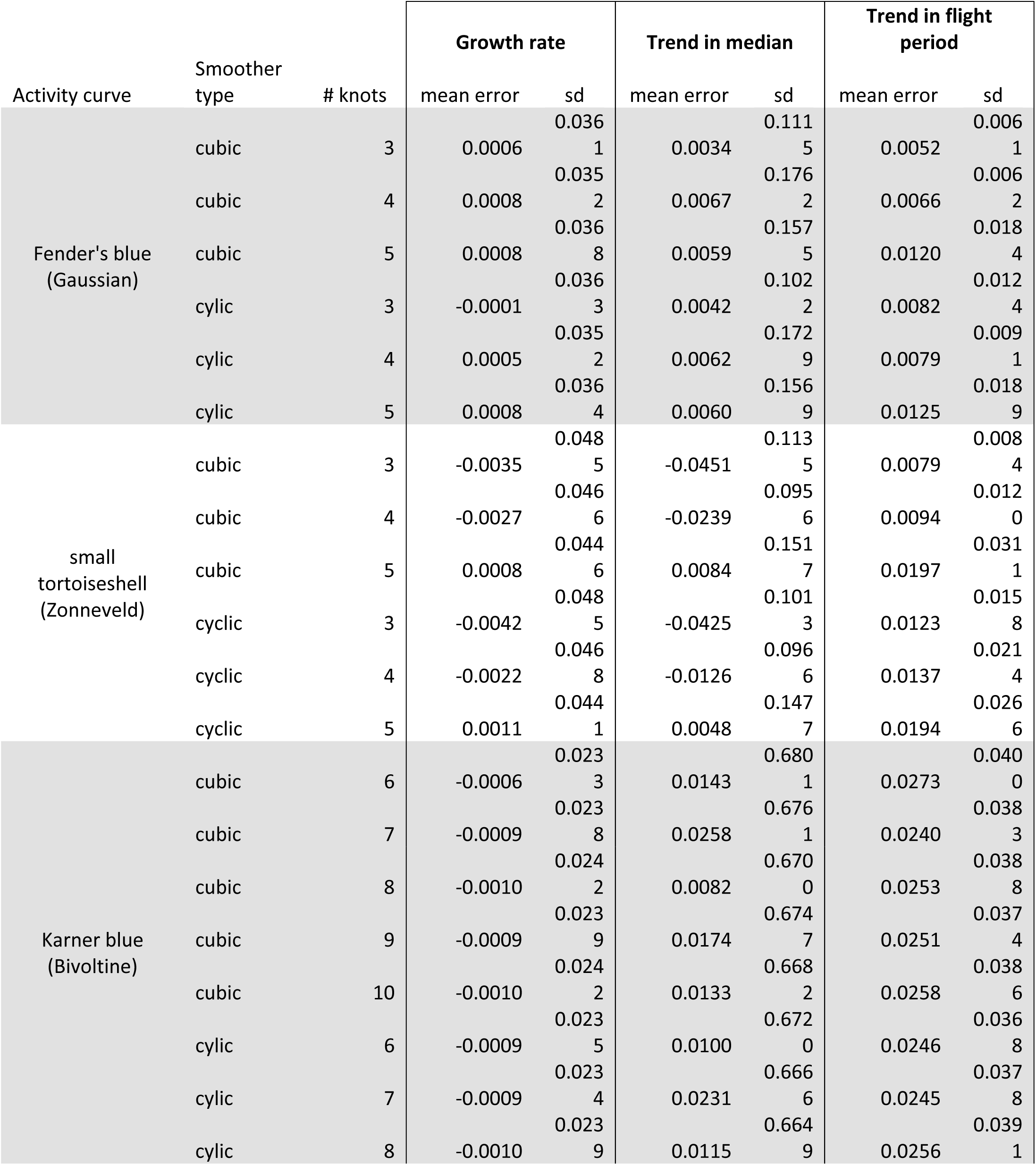
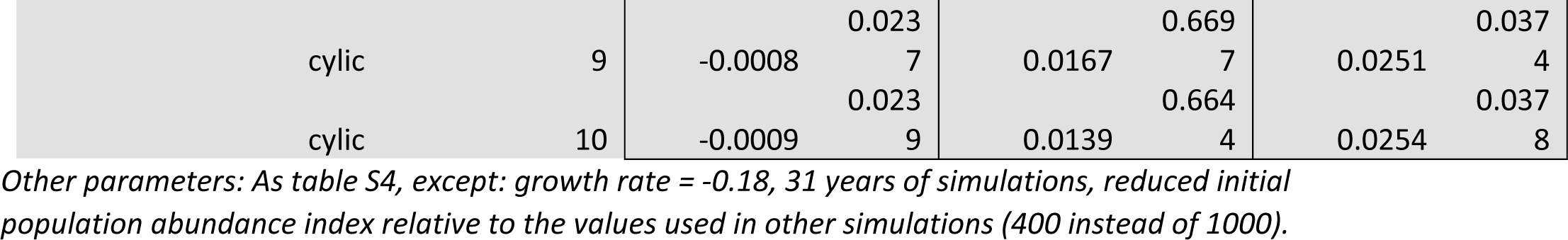
Average error (bias) and standard deviation of estimated trends (variance) as in Table S3, but using less extreme declines.

### Excluding days with few day-positives

When carrying out post-hoc analysis to estimate trends across years, it can be useful to exclude years with insufficient non-zero (“day positive”) counts in order to reduce bias in estimates. Here we focus on the extreme decline scenario (growth rate = −0.18, 30 years of data) because these exclusion rules will only matter for time series with years of low/no day-positives. When estimating growth rates, it was generally best to include every year. While for the Zonneveld distribution had mean error that was the smallest when excluding years with no day positives, in practice we are unlikely to know the underlying activity distribution, so we recommend using all years. When estimating phenology trends, error was minimized when excluding years with fewer than 2 day positives (Gaussian), 1 day positive (Zonneveld), or 2 (median) or 4 (flight period) day positives (Bivoltine). Across other parameter combinations (e.g. other knot counts and anchor distances), we found that a minimum of 1 or 2 day positives performed the best for univoltine activity curves and performed reasonably well for bivoltine activity curves. For simplicity and consistency, we recommend excluding years with fewer than 2 day positives when estimating phenology trends to any activity curve.

**Table S6:**
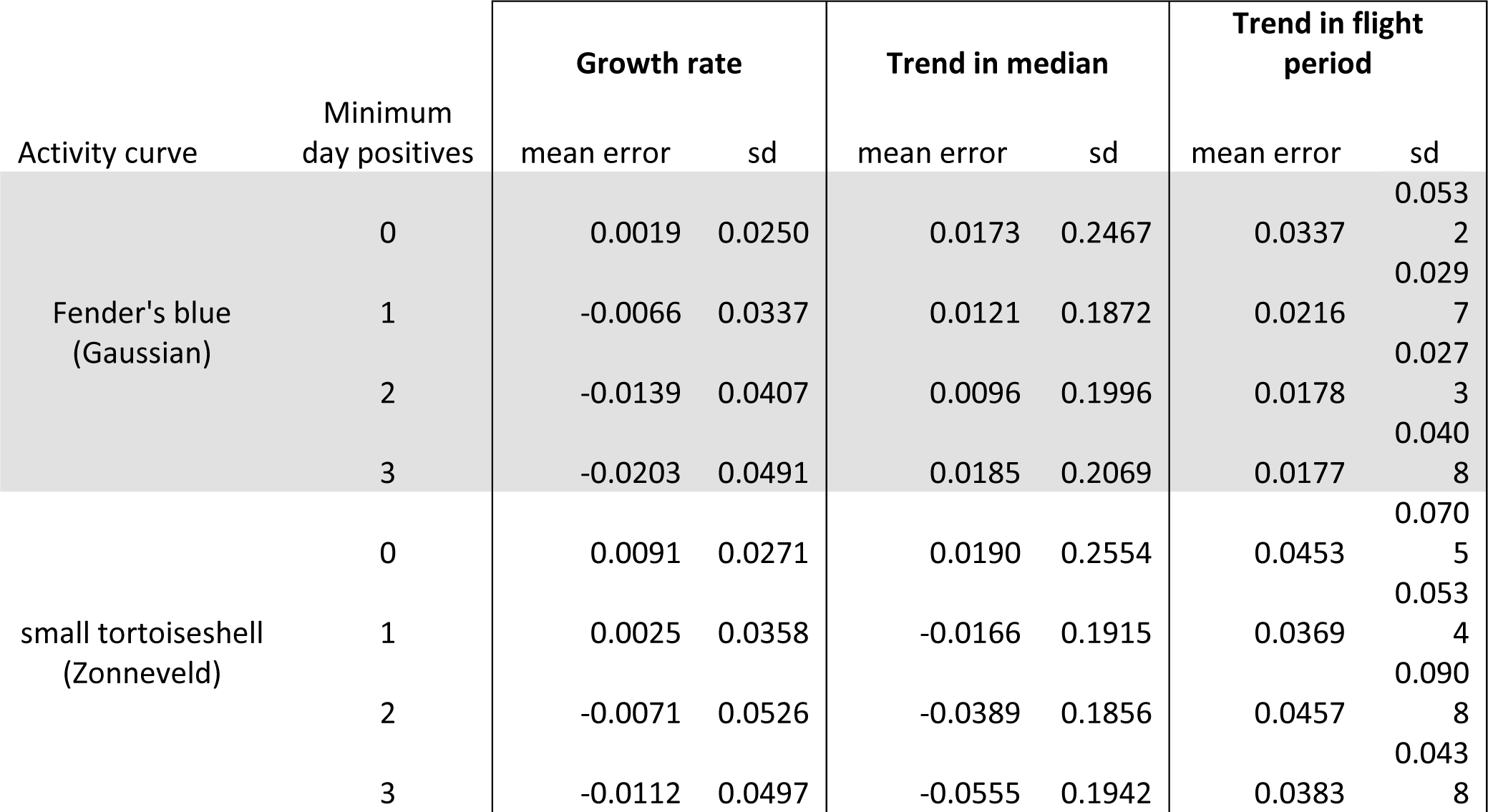
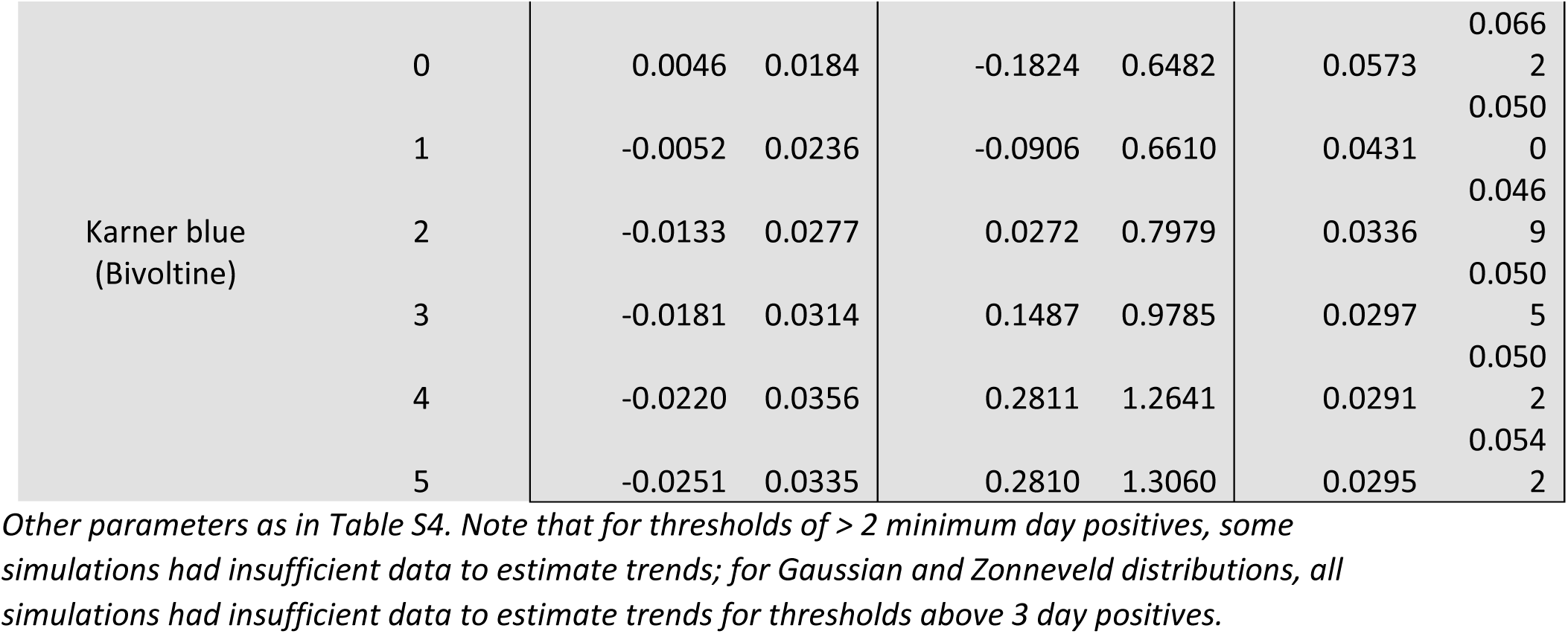
average error (bias) and standard deviation of estimated trends (variance) when excluding years with too few day positives.

